# GLIA CELLS ARE SELECTIVELY SENSITIVE TO NANOSIZED TITANIUM DIOXIDE MINERAL FORMS

**DOI:** 10.1101/2025.08.25.671597

**Authors:** Eszter Geiszelhardt, Erika Tóth, Károly Bóka, Katalin Schlett, Norbert Bencsik, Krisztián Tárnok

**Author notes:** **Author Contributions:** Conceptualization, K.T. and K.S.; methodology, K.T., K.S. and E.T.; investigation, E. G.; K.T.; writing—original draft preparation, E.G. and K.T.; writing—review and editing, E.G., E.T., N.B., K.T. and K.S; visualization, E.G.; supervision, K.S. and K.T.; project administration, N.B. and K.T.; funding acquisition, E.T., N.B. and K.S. All authors have read and agreed to the published version of the manuscript.

## Abstract

Nanosized titanium dioxide is widely used by the industry e.g. in pigments, suncreams and food colors. Its environmental and biological effects have been investigated in the past, however, few studies have focused on its crystal structure-specific effects. In our experiments, the toxicity of two types of nanoparticles was examined on primary neural cultures with different cell-compositions, using MTT and LDH assays. Primary murine cell cultures containing only astroglia cells originated from two brain regions, as well as mixed neurons and glia cells or microglia cells exclusively, were treated with anatase and rutile TiO_2_ nanoparticles at varying concentrations for 24 or 48 hours. Our results show that neither anatase nor rutile nanoparticles reduced viability in cell cultures containing a mixture of neurons and glial cells, independently of the applied concentration and treatment time. Rutile but not anatase form induced cell death in cortical astroglia cultures already at 24 hours of treatment above 10 µg/mL, while hippocampus-derived glial cultures were much less sensitive to rutile. The rutile form also damaged microglia. These findings suggest that products containing rutile-form nano-titanium particles may pose a targeted risk to astroglia and microglial cells in the central nervous system.

## 1. Introduction

Titanium dioxide (titanium (IV) oxide, TiO_2_) is a naturally occurring oxide of titanium that is found as a mineral in magmatic rocks and hydrothermal veins. It naturally occurs in different crystal structures. The most prevalent of these are anatase, brookite and rutile. Rutile is considered the most stable form of TiO_2_, while anatase and brookite are classified as metastable. The brookite form is the most unstable of the three mentioned above, and its artificial production is the most challenging (Rahimi et al., 2016).

Titanium dioxide has a wide range of industrial applications: around 80% of the world’s titanium dioxide production is used to produce pigments (e.g. titanium white) for paints, varnishes, plastics and papers. In powder form, titanium dioxide is also used as opacifier to color several products including plastics, medicines (i.e. pills) and toothpastes (Bischoff et al., 2020). An additional 8% is used in other pigment applications, such as printing inks, and it also plays a significant role in scientific research, particularly in the synthesis of new TiO₂-based composite materials (Deng et al., 2020). TiO_2_ can also be found in foodstuffs under the name of the E171 supplement, especially in ice creams, marshmallows, chewing gum, creamers etc. (Weir et al., 2012). TiO_2_ is a well-known ingredient in skin care products: it is used in sunscreens and cosmetics due to its light-scattering properties and resistance to discoloration (Racovita, 2022; Tucci et al., 2013). Titanium dioxide is typically used in the industry in the form of nanoparticles with a size range of 100 nm to 500 nm, including ultrafine particles averaging between 10 and 50 nm (Baranowska-Wójcik et al., 2020; Peters et al., 2014). As nanoparticles, anatase has a spherical shape, while rutile resembles a rod-like structure (Wilson et al., 2015).

Until 2006, titanium dioxide (TiO_2_) was registered as a harmless substance with no known health implications (Jovanović, 2015; Kischkewitz, 2006), even though an increasing number of studies have focused on the effects of its nanoparticles on human health (Song et al., 2015, 2016). The toxicity of nanoparticles can be transmitted through various routes, including inhalation, ingestion, or injection, with each pathway exhibiting distinct characteristics (Johnston et al., 2009; Lin et al., 2020). The gastrointestinal absorption of TiO₂ nanoparticles was indirectly evaluated by analyzing titanium levels in various organs. Titanium dioxide has been detected in the blood, brain, and the small and large intestines (Heidari et al., 2019; Hu et al., 2016; Onishchenko et al., 2012; Yang et al., 2017). Moreover, several studies have shown that ingested TiO_2_ can cause a variety of problems, including inflammation or cell death, and can also affect the immune- and nervous systems (Czajka et al., 2015; Rashid et al., 2021). Due to its potential adverse health effects, foods containing more than 1% TiO_2_ by weight of the product are now banned in the United States (Baranowska-Wójcik et al., 2020). The European Union has also banned the use of TiO_2_ (E171) in foods in January 2022 (European Commission, 2022).

The neurotoxic potential of TiO₂ nanoparticles has been investigated in several *in vivo* and *in vitro* studies, with particular focus on their ability to cross the blood-brain barrier (Song et al., 2015). For example, studies using both *in vitro* and *in vivo* models have shown that TiO₂ nanoparticles can exert toxic effects on rat neuronal cells, including reduced neuroblast proliferation and activation of microglia (Valentini et al., 2018). Moreover, it was reported that administering TiO_2_ nanoparticles at different doses can reduce the number of tyrosine hydroxylase-positive neurons (Heidari et al., 2019). Mechanistically, TiO₂ nanoparticles have been shown to disrupt the ubiquitin–proteasome system and associated protein degradation pathways in PC12 cells, leading to increased cytoplasmic α-synuclein accumulation and pathological aggregation (Wu & Xie, 2016). Based on the chemical characteristics of TiO₂, the most plausible toxic effects are related to oxidative stress and/or the induction of neuroinflammation (Hou et al., 2019; Song et al., 2016). TiO₂ nanoparticles also exert harmful physiological effects on other brain cell types, including the induction of oxidative stress in glial cells (Huerta-García et al., 2014). This is particularly noteworthy, as neuronal damage can be a secondary consequence of primary glial dysfunction (Heneka et al., 2010), given that astrocytes play a central role in protecting against oxidative stress (Y. Chen et al., 2020). However, astrocytes from different brain regions exhibit distinct gene expression profiles, metabolic activity, and inflammatory responses (Batiuk et al., 2020; Endo et al., 2022; Morel et al., 2017). Therefore, TiO₂-induced toxicity, oxidative stress, or immune activation may vary significantly depending on the origin of glial cells.

Studies have rarely focused on the specific effects of different crystal forms - anatase and rutile - in detail. Therefore, the aim of this study is to investigate the neurotoxic effects of different crystal forms of TiO₂ ultrafine nanoparticles (< 50 nm) on various murine primary neural cultures; including glial cells of hippocampal and cortical origin. We also examined the concentration-dependent effects on the viability of cultures containing both neurons and glia (mixed cultures) or pure microglia. The potential impact of varying treatment times (24 or 48 hours) on nanoparticle-evoked responses was also investigated. We hope that delineating cell-type specific responses will contribute to understanding the risk of TiO_2_ load on specific cell types of the CNS.

## 2. Materials and Methods

### 2.1. Animal handling

For the experiments CD1 wild-type mice (Charles River Laboratories, Wilmington MA, USA; organism: RRID:IMSR_CRL:22) were used as cell donors. The animals were housed at 22±1 °C with 12/12 -hour dark/light cycle. Water and food were available ad libitum. Experiments were carried out in accordance with the Hungarian Act of Animal Care and Experimentation (1998, XXVIII) and with the directive 2010/63/EU of the European Parliament and of the Council of 22 September 2010 on the protection of animals used for scientific purposes (“Directive 2010/63/EU of The European Parliament and of the Council of 22 September 2010 on the Protection of Animals Used for Scientific Purposes,” 2010). Experimental protocols were approved by the Animal Care and Use Committee of Eötvös Loránd University. Sacrifying the animals to obtain brain tissue and prepare primary neuronal cultures is regarded as organ donation. All possible efforts were made to minimize the number of animals used.

### 2.2. Gene expression analyses using public datasets

For gene expression analyses we used the GSE198024 bulk RNA-seq, normalized expression dataset published by Endo et al (2022) (Endo et al., 2022), available at the Gene Expression Omnibus (GEO). All analyses were performed using the web-based bioinformatic platform useGalaxy (v25; http://usegalaxy.eu). From the original dataset, the hippocampal (HIP) and cortical (CX) samples including motor-, sensory- and visual cortices (MCX, SCX and VCX, respectively) were selected with gene expression values derived from four biological replicates. To ensure reproducibility of the results, the original sample identifiers were retained. Expression values were log_2_-transformed prior to statistical analysis. Differential gene expressions were assessed using the edgeR package (v3.36), which included filtering of low-expression genes (CPM < 1), dispersion estimation, and exact test-based comparison. Genes with a false discovery rate (FDR) < 0.05 and absolute log_2_ fold change (|log2FC|) > 1 were considered significantly differentially expressed. Heatmaps were generated using the heatmap2 function (v3.2.0) with row-based Z-score normalization. Principal component analysis (PCA) was performed using the prcomp function in R and visualized via ggplot2 (PCA plot w ggplot2 v3.4.0). For molecular signature analyses gene sets from GSEA Molecular signature database (https://www.gsea-msigdb.org) were used (see Supplementary **Table S1**).

### 2.3. Preparation of cell cultures

Experiments were performed on different types of primary neural cell cultures. (i) Mixed cultures containing both neurons and glia cells, as well as cultures containing (ii) only neurons, (iii) hippocampal astroglia, (iv) cortical astroglia or (v) microglia were prepared. Cellular composition of the cultures was routinely verified by anti-IIIβ-tubulin, anti-GFAP, and anti-Iba1 immunostaining, as described previously (Szentgyörgyi et al., 2025).

#### 2.3.1. Preparation of mixed neuron and astroglia and pure neuronal cultures

Mixed (where both neurons and astroglia cells were present) and pure neuronal cultures were prepared from 14-14.5 days old mouse embryos according to the protocol previously reported by our laboratory (Tárnok et al., 2008). Briefly, the brains of the animals were isolated under aseptic conditions, the meninges were removed, the cortices were dissected and incubated in 0.5 mg/mL trypsin-EDTA (Thermo Scientific, Hungary) solution at 37 °C for 15 min. Cells were seeded onto poly-L-lysine (Sigma-Aldrich-Merck, Hungary) coated 96-well tissue culture plates (Greiner Bio-One, Hungary) at 8×10^4^ cells/well density. For microscopy, 1×10^5^ cells were plated onto poly-L-lysine – laminin (Sigma-Aldrich-Merck, Hungary) coated glass coverslips in 24-well plates. Cells were cultivated at 37 °C in 5% CO_2_ / 95% air atmosphere in Neurobasal Plus medium supplemented with 5% FCS (Neurobasal Plus, 2% B27, 0.5 mM Glutamax, 5% FCS, 40 µg/mL gentamycin, 2.5 µg/mL Amphotericin B, Thermo Scientific, Hungary).

In cortical mixed cultures, complete medium change with the plating medium was performed on the first day after preparation (days *in vitro*, DIV1). On day 4 (DIV4), half of the culture medium was replaced with fresh, FCS-free medium (Neurobasal Plus, 2% B27, 0.5 mM Glutamax, 40 µg/mL gentamycin, 2.5 µg/mL Amphotericin B, Thermo Scientific, Hungary). Pure neuronal cultures were cultivated similarly but 5% FCS was present only until DIV1 and cells were treated with cytosine– arabinofuranoside (CAR, 10 μM; Sigma-Aldrich-Merck, Hungary) 24 h after plating to prevent the further division of non-neuronal cells. Titanium dioxide-treatments were performed on the 7^th^ day of cultivation.

#### 2.3.2. Preparation of pure astroglia cultures

Primary cortical astroglia cells were obtained from 1-3 day-old (P1-P3) neonatal CD1 pups according to the protocol used in our laboratory (Tárnok et al., 2010). Briefly, the brains were aseptically isolated, the meninges, the cerebellum and olfactory bulbs were removed, and the cortexes were dissected and incubated in trypsin-EDTA solution (0.5 mg/mL, Thermo Scientific, Hungary) containing 0.05% DNase (Sigma-Aldrich-Merck, Hungary) for 15 min at 37 °C. Cells obtained from 3 hemispheres were plated per 10 cm Petri dish, surface-treated with poly-L-lysine (Sigma-Aldrich-Merck, Hungary). Cultures were grown in HDMEM medium containing 10% FCS (high glucose DMEM, 10% FCS, 40 µg/mL gentamycin, 2.5 µg/mL Amphotericin B, Sigma-Aldrich-Merck, Hungary), with a complete medium change in every 4 days. For the titanium dioxide treatments, cells were plated into poly-L-lysine coated 96-well tissue culture plates at a cell density of 2.5×10^4^ cells/well or onto poly-L-lysine – laminin (Sigma-Aldrich-Merck, Hungary) coated glass coverslips in 24-well plates at a cell density of 1×10^5^ cells/well. TiO_2_ treatments were performed on the 2^nd^ day after plating.

The hippocampal astroglia cultures were obtained from 16-day-old (E16) CD1 mouse embryos. Hippocampi were cut from the cleared cortices, and cells were isolated by trituration after digestion (15 min, 37°C) with trypsin-EDTA solution (0.5 mg/mL, Thermo Scientific, Hungary) containing 0.05% DNase (Sigma-Aldrich-Merck, Hungary) and then plated in 10 cm Petri dishes surface-treated with poly-L-lysine. Cultures were maintained in the same manner as cortical, neonatal astroglia cultures. For the treatments, cells were plated in a 96-well tissue culture plate coated with poly-L-lysine (Sigma-Aldrich-Merck, Hungary) at a cell density of 2.5×10^4^ cells/well. Treatments were performed on the 2^nd^ day after plating.

#### 2.3.3. Preparation of microglial cultures

Microglia cultures were prepared from neonatal astroglial cultures. After 1-2 weeks, microglia cells appeared on the surface of the confluent astroglia cells. Microglial cells were washed off from the surface of the astroglia cells with HDMEM (high glucose DMEM, 10% FCS, 40 µg/mL gentamycin, 2.5 µg/mL Amphotericin B, Sigma-Aldrich-Merck, Hungary) and were plated on 96-well plates surface-treated with poly-L-lysine (Sigma-Aldrich-Merck, Hungary) at a cell/well ratio of 2.5×10^4^ cells. Microglial cells were treated with titanium dioxide one day after plating.

### 2.4. Preparation, application and verification of titanium dioxide solutions

Aqueous dispersions of anatase and rutile form TiO_2_ nanoparticles (NP’s) were provided by Holikem Ltd. (Hungary). Anatase dispersion contained 5.5 wt% of spherical TiO_2_ particles, while the rutile dispersion consisted of 13.17 wt% rod-shaped TiO_2_ particles. The structure of anatase and rutile nanoparticles was verified by transmission electron microscopy (TEM, **Figure 1**). For TEM preparation, nanoparticles dissolved in distilled water were dropped onto a carbon-coated Formvar grid and negatively stained according to Tóth et al. (2017) (Tóth et al., 2017). After air drying, the grids were examined by a Hitachi 7100 TEM. Morphometric analysis of calibrated electron microscopic images was performed using ImageJ/Fiji v1.54p (Schindelin et al., 2012), measuring the length and diameter of 30 NP’s per sample. Data were visualized as individual values with medians using Prism v10 (GraphPad, USA).

**Figure 1.**
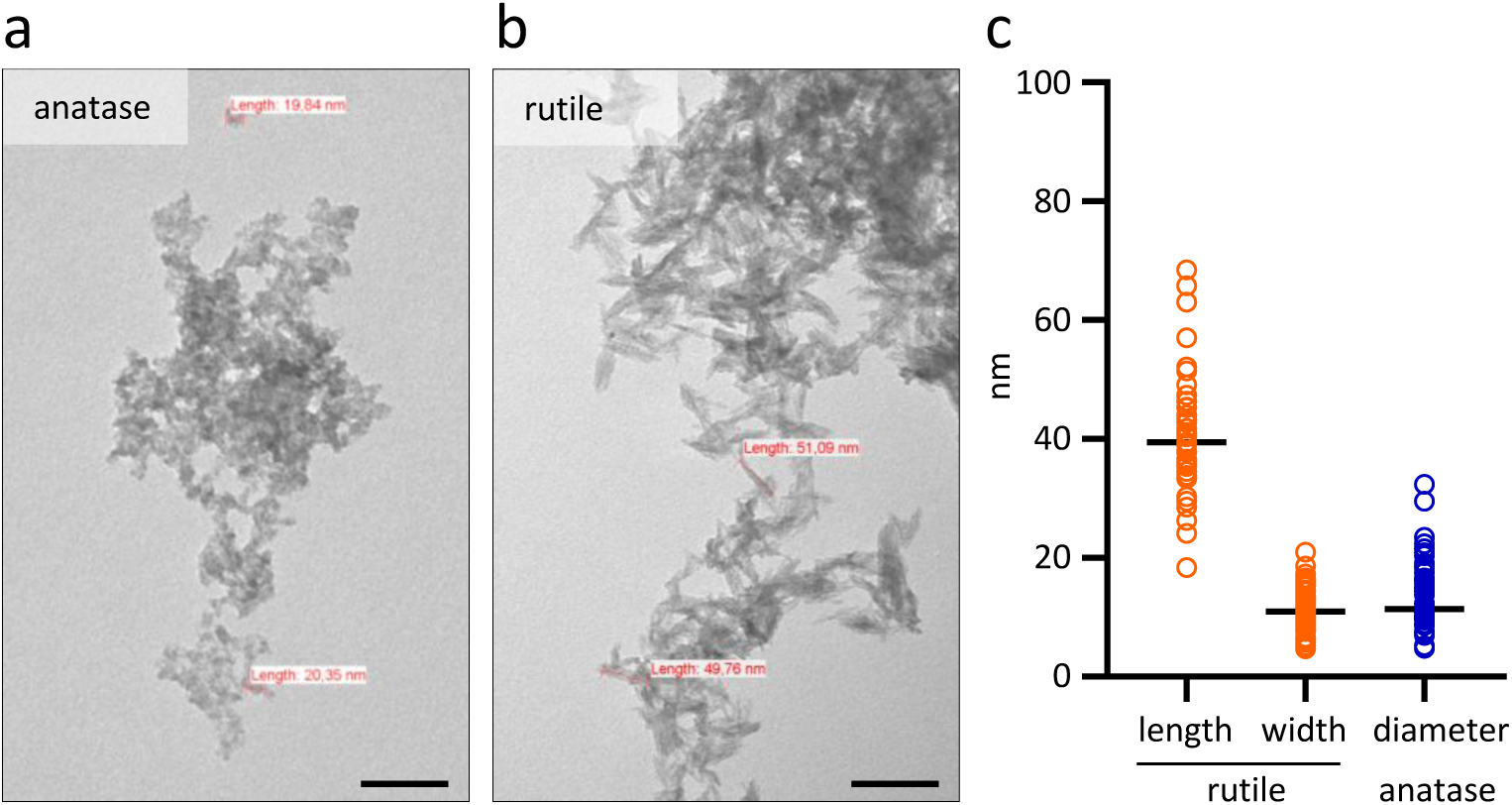
Structure of anatase (a) and rutile (b) nanoparticles. Anatase has a spherical shape with a 15.8±1.7 nm average diameter, while rutile has a rod-like shape, with a 46.7±2.2 nm average length and 13.7±0.7 nm average diameter. The scalebar indicates 100 nm. (**c**) Morphometric analysis of crystal length and diameter. *n*=30

For the experiments, dispersions were prepared by diluting the stock of TiO_2_ nanoparticles in distilled water, phosphate buffered saline (PBS) and Neurobasal Plus (Thermo Scientific, Hungary) medium to obtain a 10 mg/mL stock solution. To achieve a homogenous dispersion of the nanoparticles, the stock solutions were sonicated at 50 W for at least 30 minutes. No differences were observed between the solvents; titanium dioxide was evenly dispersed in all media.

### 2.5. Viability measurement using MTT method and cytotoxicity measurement using LDH assay

The effects of titanium dioxide on cell viability were assessed using the MTT (3-(4,5-dimethylthiazol-2-yl)-2,5-diphenyl tetrazolium bromide, Sigma-Aldrich-Merck, Hungary) reduction assay (Mosmann, 1983). MTT was applied at a final concentration of 200 μg/mL to the cells grown in 96-well plates, and plates were incubated for 20-35 min at 37 °C in a tissue culture incubator. The cells and formazan crystals were dissolved in acidic (0.08 M HCl) isopropyl alcohol (Sigma-Aldrich-Merck, Hungary). The optical density (OD) was determined with a Multiskan EX spectrophotometer (Thermo Scientific, Hungary) at 570 nm, with reference wavelength of 620 nm. Because titanium dioxide particles may interfere with spectrophotometric measurements, the treatment was also performed on cell-free wells in all applied titanium dioxide concentrations. These values, as OD backgrounds, were subtracted from the ODs obtained from cell-containing wells. Viability data were presented as a percentage of the untreated (without nanoparticles) control wells (relative cell viability %).

LDH (lactate dehydrogenase) enzyme activity assay (CyQuant LDH Cytotoxicity Assay, C20300; Thermo Scientific, Hungary) was used to determine the cytotoxicity of the NPs’ treatments. Equal volumes of the collected supernatants were pipetted into a new microtiter plate and treated according to the manufacturer’s instructions. Optical density values were determined using a Multiskan EX ELISA reader (Thermo Scientific, Hungary) spectrophotometer at 492 and 620 nm wavelength. For the visualization of datasets, see the ‘Datasets, statistics and data visualization’ section. LDH-assay was applied in microglia and pure cortical astroglia cultures, where the viability-reducing effect of titanium dioxide was evident in the MTT measurements.

### 2.6. Datasets, statistics and data visualisation

For MTT measurements of both astroglia and mixed cultures as well as cortical glial cultures, 5 independent experiments were performed using anatase and rutile nanoparticles with treatment times of 24 and 48 h, with 6-8 parallel measurements per treatment. For hippocampal astroglia cultures, results were obtained from 2 independent experiments with 6-8 parallel measurements per treatment. Data for microglia cultures were obtained from 6-8 parallels per treatment.

LDH measurements were carried out on cortical astroglia and microglia cultures. In cortical glia cultures, 2 independent measurements with 4-6 parallels per treatment, in microglia cultures, one measurement with 3-6 parallels per treatment were performed.

MTT results (OD values obtained after subtracting the background measured in TiO_2_-only containing wells with identical titanium dioxide load) were normalized to the control ODs (solvent treated cultures) and presented as a percentage (relative cell viability %). Parallel measurements within a plate were considered as a single dataset. LDH data were calculated using the following formula: cytotoxicity % = [(LDH release measured on treated well - spontaneous LDH release) / (maximum LDH release - spontaneous LDH release)]×100.

Statistical analysis was performed with Jamovi (v2.6.26, jamovi.org). Normal distribution of data was tested by Shapiro-Wilk test at p<0.05 significance level. When the data showed a normal distribution, one-way analysis of variance (ANOVA) was performed, using Tukey’s test as post-hoc test. If the data were not normally distributed, non-parametric Kruskal-Wallis test with Dwass-Steel-Critchlow-Fligner pairwise comparison was chosen. Significant differences compared to control values were indicated, using the symbol * (p<0.05: *; p<0.01: **; p<0.001: ***).

The comparison of anatase- and rutile-induced effects in 24 and 48 h treatments were performed using Student-t test. Normal distribution of data was tested by Shapiro-Wilk test at p<0.05 significance level. If the distribution wasn’t normal, Mann-Whitney U test was used, the “$” indicates the levels significance (p<0.05: $; p<0.01: $$; p<0.001: $$$).

Data were plotted using Microsoft Excel and presented as mean ± SEM. To facilitate comparison, the datasets were presented in two ways: one graph displays the grouped data for each treatment time by the applied concentration of both crystal structures, while the second graph compares the effects of different treatment durations for a given type and concentration of nanoparticles.

## 3. Results

### 3.1. Characterization of anatase and rutile nanoparticles

TiO_2_ ultrafine nanoparticles in anatase and rutile crystal forms were characterized in aqueous dispersions by transmission electron microscopy and subsequent image analysis. Anatase nanoparticles showed isometric spherical shape (**Figure 1a**) with 15.8±1.7 nm average diameter (**Figure 1c**). TiO_2_ nanoparticles in rutile form showed anisometric, elongated, rod-like shape (**Figure 1b**) with 46.7±2.2 nm length and 13.7±0.7 nm diameter (**Figure 1c**). The dispersions were 100% pure, no other crystal formations were observed.

### 3.2. Effect of TiO_2_ nanoparticles on the viability of hippocampal astroglia cultures

The presence of TiO_2_ nanoparticles has been shown to exert a harmful effect on neural tissue, through both direct (neuronal-based) and indirect (astrocyte and microglial-mediated) pathways (Song et al., 2016). Astroglial cells have long been regarded as a “homogeneous” cell type which, in contrast to neurons, lack substantial morphological or functional diversity (Verkhratsky & Nedergaard, 2018). However, recent studies, primarily employing astrocyte-specific bulk- and single-cell RNA sequencing (scRNA-seq) techniques, have revealed that the gene expression profiles of glial cells differ significantly across brain regions (Batiuk et al., 2020; Endo et al., 2022). These studies have mainly focused on identifying distinct astrocyte subpopulations and have not specifically addressed oxidative processes (Batiuk et al., 2020; Dallérac et al., 2018; Endo et al., 2022; Xin & Bonci, 2018). The standardized and open access sharing of these published datasets has made it possible to analyze such processes independently of the original research aims. Based on the *GSE198024* astroglia bulk-RNASeq dataset, in which Endo et al. performed RiboTag-based RNA profiling on astrocytes originating from different brain regions of *Aldh1l1*-CreERT2 x RiboTag adult mice (Endo et al., 2022), we compared the transcriptome of astrocytes with hippocampal (HIP) and cortical (from moto-, sensory- and visual cortex, MCX, SCX and VCX, respectively) origin. As was anticipated based on the former publication (Batiuk et al., 2020; Endo et al., 2022), hippocampal and cortical astroglia cell are separated based on 359 differentially expressed genes (see **Figure S1a, b**). When we filtered the original normalized expression dataset to the Hallmark Reactive Oxygen Species Pathway GSEA gene set (MM3895, GSEA), principal component analyses also separated the hippocampal astrocytes from the cortical ones (**Figure S1c, d**). Systematic filtering of the 359 differentially expressed genes with oxidative stress and glutathione metabolism related gene sets from GSEA Molecular Signature Database (**Table S1**) revealed 6 genes which also separated clearly the two groups (**Figure S1e, f**).

Based on the oxidative stress-related gene set-dependent separation of brain region-derived astroglia transcriptomes and the fact that astroglia cell can preserve some brain region-specific characteristics *in vitro* (Cragnolini et al., 2018; Ernsberger et al., 1990; Kipp et al., 2008), we compared the effects of TiO_2_ nanoparticles on cultured hippocampal and cortical astrocytes. To achieve that, hippocampal astroglia was prepared from the hippocampus of 16-day-old mouse embryos, while cortical astroglia were obtained from 1 to 3-day old newborn pups. All cultures were treated with dispersions of anatase or rutile nanoparticles at concentrations of 1, 5, 10, 50, 100, 200, 500 and 1000 µg/mL for 24 or 48 h.

The effects of TiO_2_ on cell viability were determined by MTT assay. To exclude any possible incompatibility between TiO_2_ nanoparticles and the photometric MTT method, TiO_2_ treatment and MTT measurements were also performed in cell-free wells and the obtained data were taken as background (blank) values (see the Material and Methods section for details). It is important to note that above a concentration of 100 µg/mL, the applied nanodispersion already precipitates on the surface of the cells and forms a layer (“nano snow ”, see on **Figure 6a**). Thus, although treatments in the concentration range 200-1000 µg/mL in most measurements were included, we assume a biologically relevant effect only below 100 µg/mL in order to get a complete picture of the toxicity concentration range.

In case of hippocampal astroglia cultures, neither anatase nor rutile nanodispersion had any effect on cell viability after 24 h of treatment (**Figure 2a**). Treatment for 48 h did not significantly change the effect (**Figure 2b**), although prolonged treatment with rutile above 10 µg/mL resulted in significantly lower OD values compared to 24 h treatment and at 1000 µg/mL compared to the control (**Figure 2b, d**).

**Figure 2.**
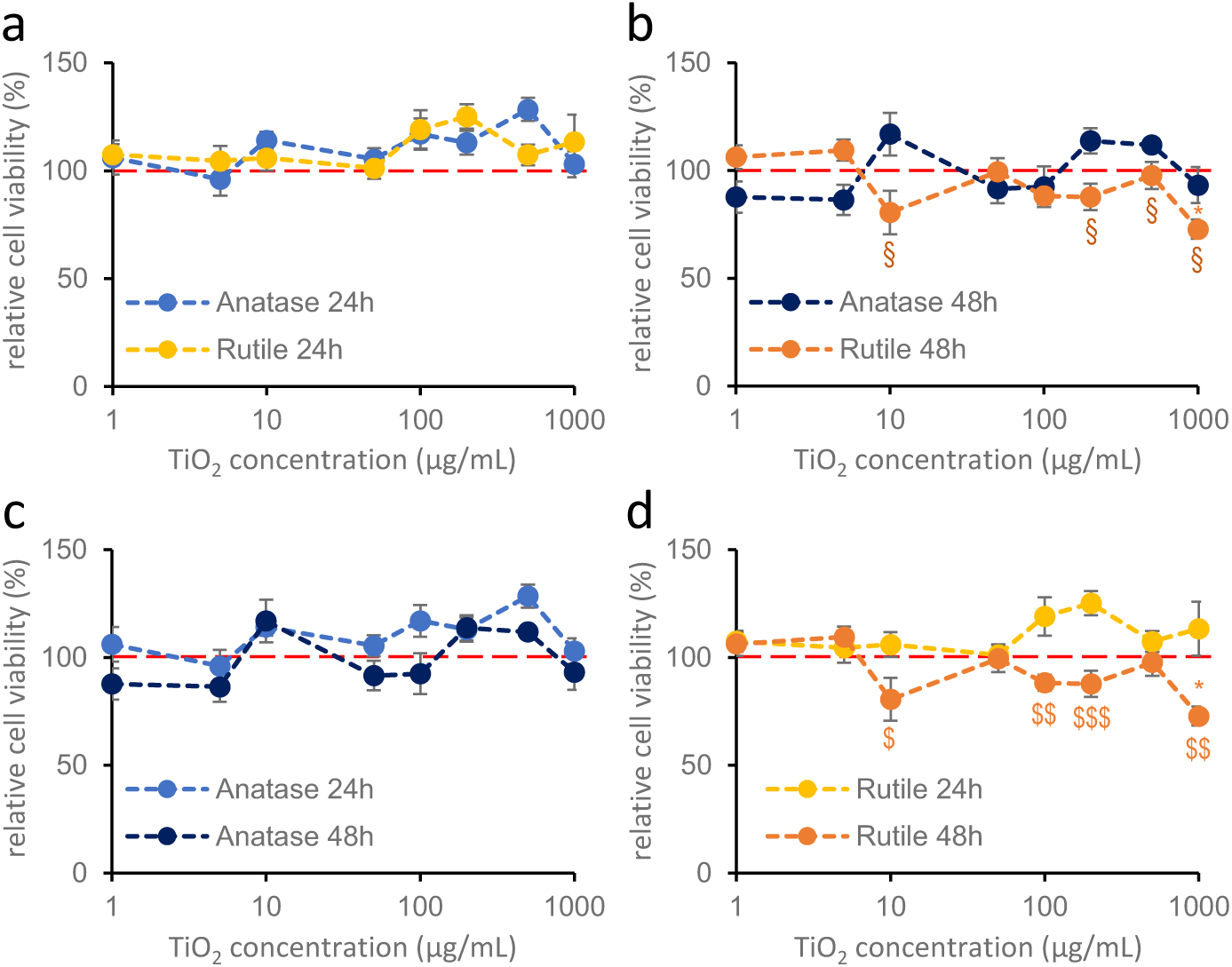
Effects of anatase and rutile on the viability of hippocampal glia cultures after 24 h (a) and 48 h (b) of treatment. Primary hippocampal glia cultures were treated with 1, 5, 10, 50, 100, 200, 500, 1000 µg/mL titanium dioxide. Blue colors indicate anatase, yellow colors indicate rutile crystal forms. Results are expressed as mean±SEM as a percentage of the untreated control. The red dotted line indicates 100% control value. § indicates comparison of anatase and rutile data (Kruskal-Wallis: p<0.05:*, §), * indicates difference from control. (**c,d**): 24- and 48-hours treatments of anatase and rutile are compared. $ indicates a significant difference between 24-48 h (**c,d**). (Student’s t test: p<0.05: $; p<0.01: $$; p<0.001: $$$) Data are derived from two independent experiments, each including 6-8 parallel measurements per crystal structure at each concentration. *n*=12-16

### 3.3. Effect of TiO_2_ nanoparticles on cortical astroglia cultures

In case of primary cortical glial cultures prepared from 1-2-day-old neonatal mice (**Figure 3**), titanium dioxide nanoparticles exerted a crystal structure-dependent effect on cell viability. Similarly to hippocampal astroglial cultures, anatase nanoparticles did not modify cell viability after either 24 or 48 h of treatment at any concentrations (**Figure 3a, b, c**). However, rutile nanodispersion significantly reduced cell viability at concentrations above 10 µg/mL (**Figure 3a, b, d**) and this effect was enhanced by longer treatment duration (**Figure 3b, d**).

**Figure 3.**
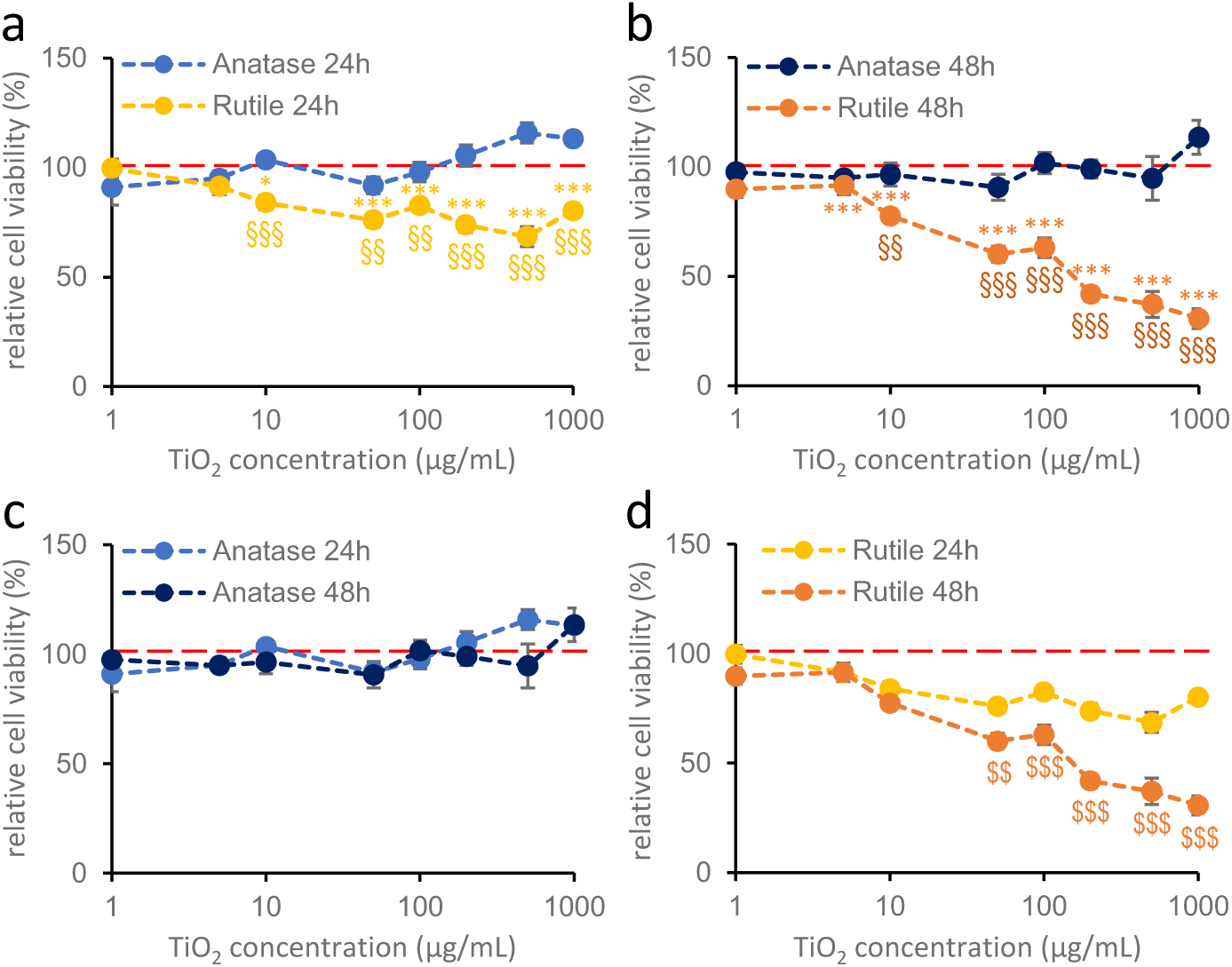
Effects of anatase and rutile on the cell viability of cortical glia cultures after 24 h (a) and 48 h (b) of treatment (MTT measurement). Primary cortical glia cultures were treated with 1, 5, 10, 50, 100, 200, 500, 1000 µg/mL titanium dioxide nanoparticles. Blue colors indicate anatase, yellow colors indicate rutile. Results are expressed as mean±SEM as a percentage of the untreated control. Data are from 5 independent measurements, with 6-8 parallel measurements per crystal structure in each concentration. The red dotted line indicates 100% as the control value. § indicates comparison of anatase and rutile, * indicates difference from control (**a,b**; Kruskal-Wallis test: p<0.05: *; p<0.01: §§; p<0.001: §§§, ***). (**c,d**): 24- and 48-hours treatments of anatase and rutile are compared, where $ indicates a significant difference between 24- 48 h (**c,d**). (Student’s t test: p<0.01: $$; p<0.001: $$$).

The MTT method is influenced by several factors, including the number of cells in the culture, the amount of available enzymes in the cells, the level of NAD(H) as a reducing agent in the reaction, and even the number of mitochondria. To confirm that the observed decrease in optical density from the MTT assay was indeed due to the induced cytotoxicity, lactate dehydrogenase activity (LDH method) was also determined in the extracellular environment as an alternative measure of cell viability. Due to the “nano snow” phenomenon described in the previous section, we tested only the 1-100 µg/mL concentration range in these experiments. Treatments were conducted for 24 and 48 h, as well (**Figure 4**). LDH measurements showed partially similar results as the MTT assays. After 24 h, no increase in cytotoxicity was observed in case of anatase, but a significant increase was detected when rutile nanoparticles were present at concentrations starting from 10 µg/mL (**Figure 4a**). This supports the decrease in cell viability observed in the MTT assay. However, in contrast to the MTT results, a significant increase in cytotoxicity was observed for anatase nanodispersion at concentrations above 50 µg/mL after 48 hours (**Figure 4b, c**). For rutile nanodispersion, the increase in cytotoxicity was similar what was observed in the MTT assay, although a statistically significant difference was detected only at 100 µg/mL (**Figure 4b, d**). In both cases, the effect was enhanced by longer treatment durations (**Figure 4c, d**).

**Figure 4.**
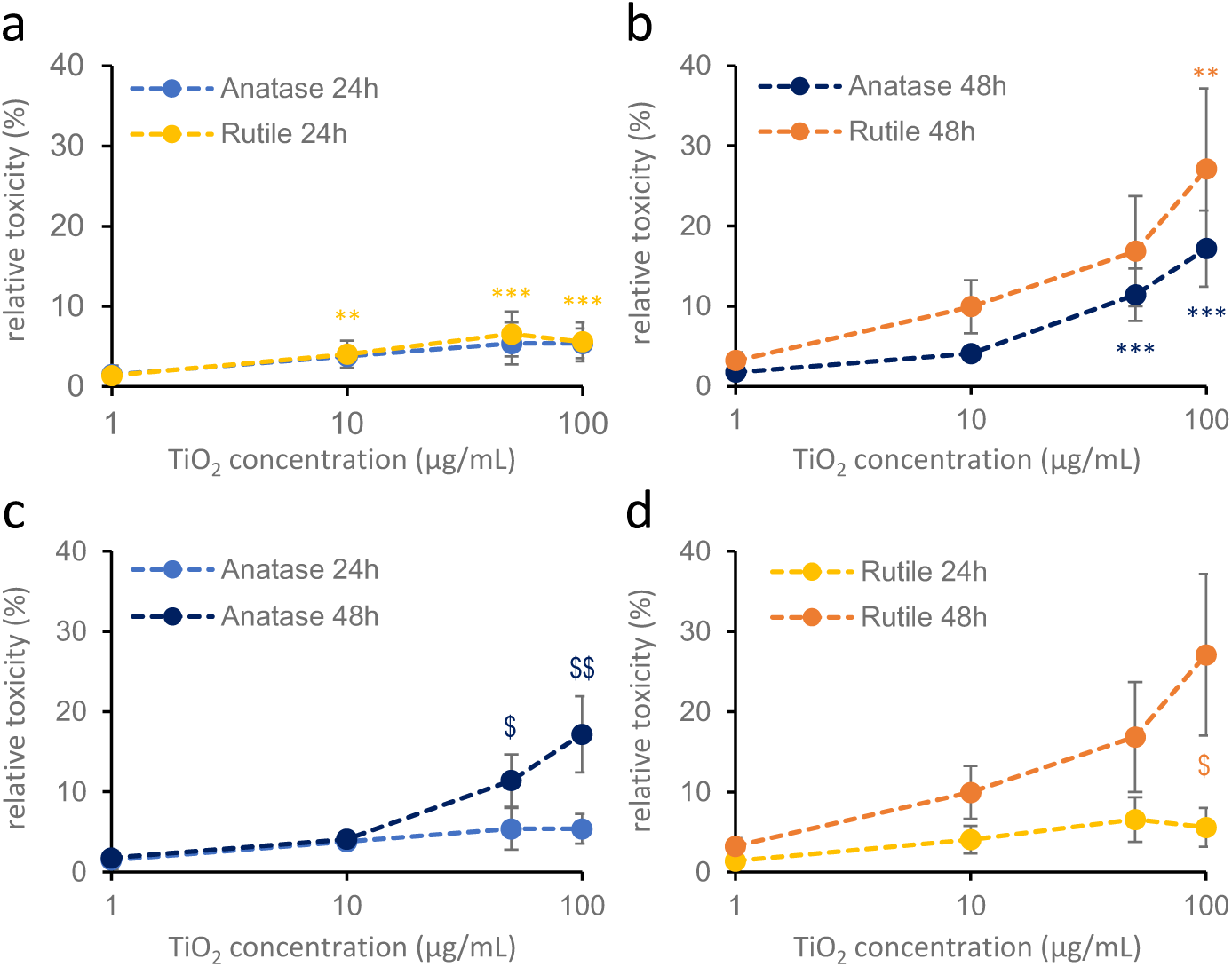
Cytotoxic effect of 24 h (a) and 48 h (b) anatase and rutile nanodispersion treatment on cortical astroglial cells. (LDH measurement). Primary cortical astroglial cell cultures were treated with 1, 10, 50, 100 µg/mL titanium dioxide. Blue colors indicate anatase, yellow colors indicate rutile nanoparticles. Results are expressed as mean±SEM. Data are from 2 independent measurements, 3 parallel measurements per crystal structure in each concentration. One-way ANOVA: p<0.05: *; p<0.01: **; p<0.001: ***, compared to untreated values) On figures **c** and **d**, 24 and 48 hours treatments are compared. (Student’s t test: p<0.05: §; p<0.01: §§)

### 3.4. Effects of TiO_2_ nanoparticles on primary cortical cultures containing both neurons and astroglia cells

In brain tissue, glial cells are never found in isolation but always in interaction with neurons, therefore we also established primary mixed cortical cultures comprising both astrocytes and neurons. The mixed cultures were treated on the 7^th^ day *in vitro* (DIV7), when the neurons had already formed a network over a layer of astroglia cells. As in previous experiments, cells were treated with dispersions of anatase or rutile nanoparticles at concentrations of 1, 5, 10, 50, 100, 200, 500 and 1000 µg/mL for 24 or 48 h and cell viability was determined by MTT assay.

Treatment with anatase at concentrations below 50 µg/mL, regardless of the treatment-time, did not cause significant changes compared to control wells. At concentrations between 50 and 1000 µg/mL, anatase treatment significantly increased the OD values of the MTT assay after 24 hours, but this effect was only observed at 1000 µg/mL after 48 hours (**Figure 5a, b**). In case of 24 h treatment, rutile nanodispersion significantly increased the relative viability at concentrations above 100 µg/mL (**Figure 5 a,b,d**), with a stronger effect than that of anatase. Interestingly, no effect on cell viability was observed after 48 hours of rutile treatment (**Figure 5d**).

**Figure 5.**
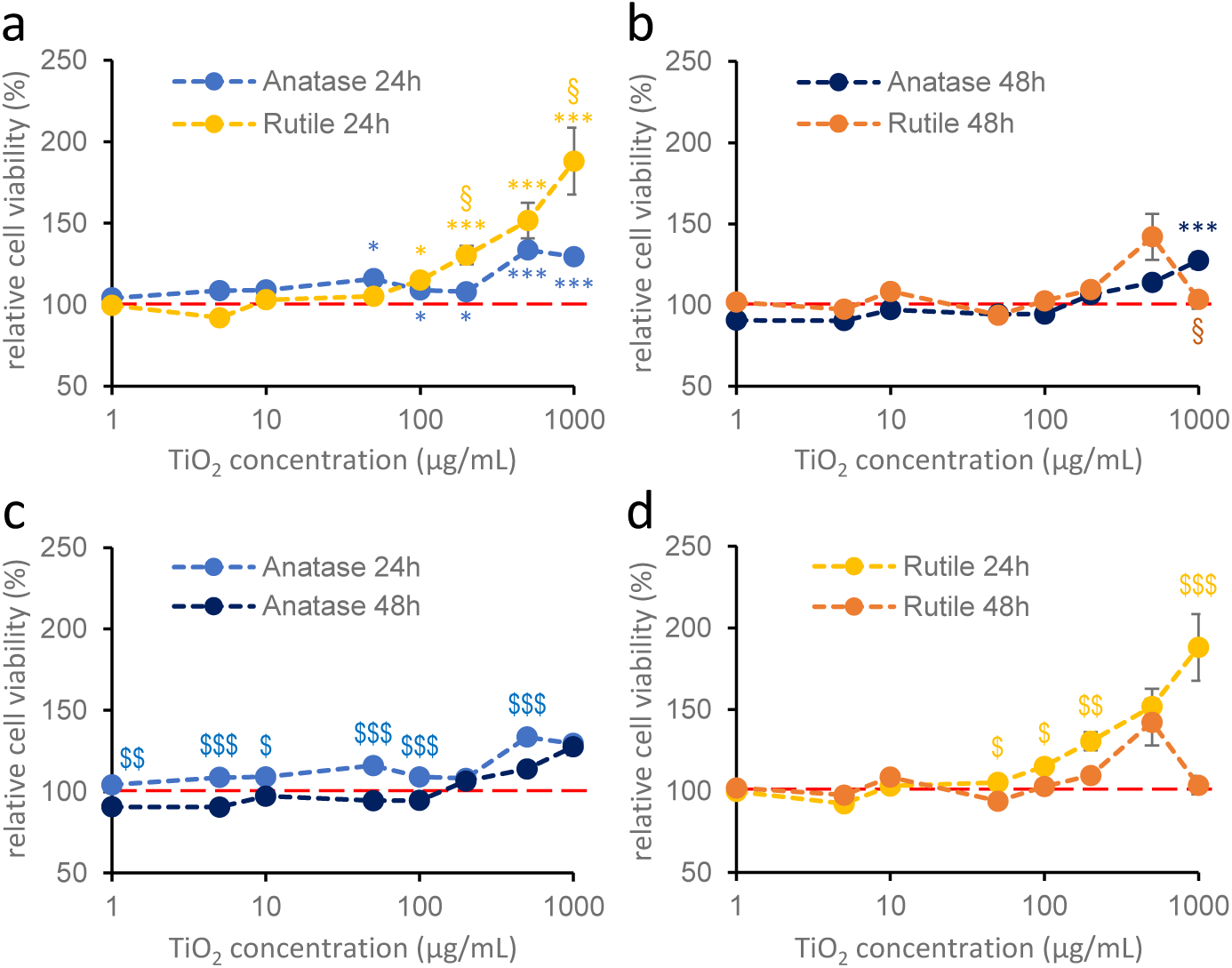
Effects of anatase and rutile on the viability of mixed cultures after 24 h (a) and 48 h (b) of treatment. Primary mixed cultures were treated on DIV7 with 1, 5, 10, 50, 100, 200, 500, 1000 µg/mL titanium dioxide. Blue colors indicate anatase, yellow colors indicate rutile. Results are expressed as mean±SEM as a percentage of the untreated control. Data are from 5 independent measurements, with 6-8 parallel measurements per crystal structure in each concentration. The red dotted line indicates 100% as the control. § indicates comparison of anatase and rutile, * indicates difference from control (a,b; Kruskal-Wallis test: p<0.05: §,*; p<0.01: **; p<0.001: ***). (**c,d**): 24- and 48-hours treatments of anatase and rutile are compared. $ indicates significant difference between 24-48 h (c,d). (Student’s t test: p<0.05: $; p<0.01: $$; p<0.001: $$$)

Our results show that none of the TiO₂ nanoforms had a viability-reducing effect on mixed neuronal-glial cultures after either 24 or 48 hours of treatment, even when very high concentrations were applied. To demonstrate that neurons themselves are not sensitive to TiO_2_ exposure, pure neuronal cultures without any supportive astroglial cells were also treated with a concentration of 100 µg/mL of both anatase and rutile nanoparticles. According to our results, anatase increased the relative viability to 121±12%, while rutile reduced it to 90±6%. However, these changes were not statistically significant compared to the control (100 ± 5%; one-way ANOVA, p = 0.093). These results suggest that TiO₂ nanoforms do not exert direct cytotoxic effects on neurons under the tested conditions.

### 3.5. Effect of TiO_2_ nanoparticles on microglial cells

Microglia, as the resident immune cells of the brain, are particularly sensitive to the presence of nanoparticles and exhibit strong phagocytic activity. Their activation plays an important role in many neuroinflammatory conditions and neurodegenerative pathologies (Hickman et al., 2018), and they are also highly sensitive to oxidative changes in their environment (Ishihara & Itoh, 2023; Rojo et al., 2014). Phagocytotic activity of these cells was clearly visible in the cultures. After 48 h of treatment with 50 µg/mL TiO_2_, the titanium dioxide “nano snow” formed on the cell-surface disappeared from the surrounding area of microglia cells (**Figure 6a**, red circles). In parallel, microglial cells changed their shape from the ramified morphology to the rounder, activated and phagocytosing form (**Figure 6a**).

**Figure 6.**
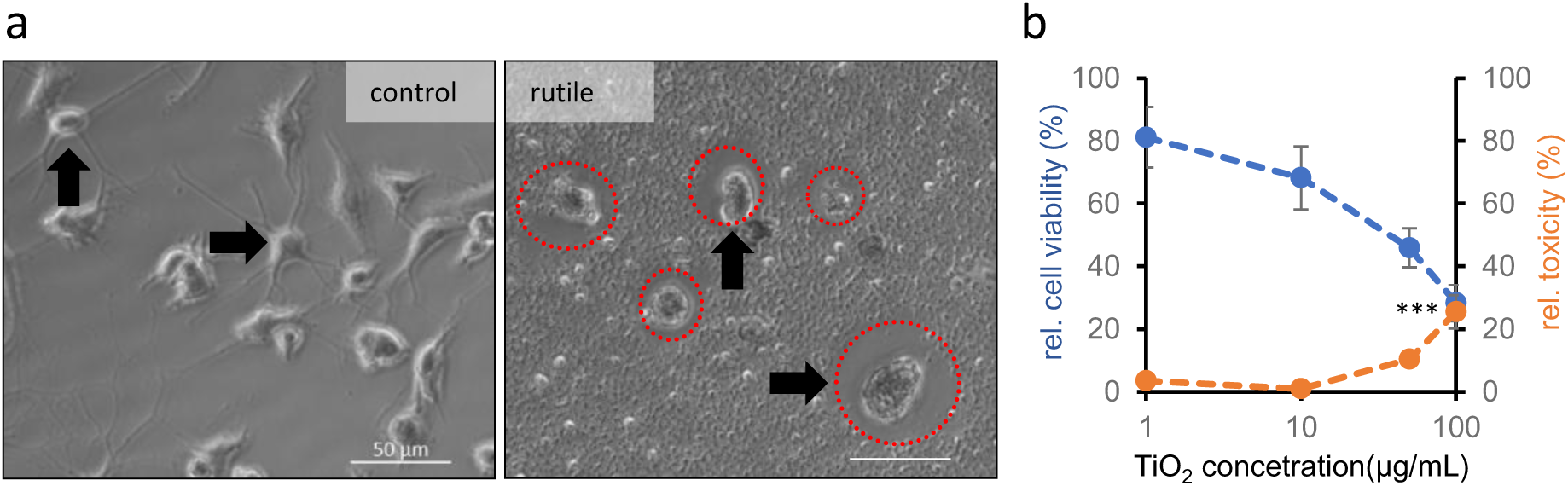
Microglia cells phagocyte TiO_2_ nanoparticles. (**a**) The shape (arrow) of cells treated with rutile (48h, 50 µg/mL, right) is changed compared to control cells (arrows on the left). The “nano snow” is not visible around the treated cells (red circles), as microglial cells likely phagocytosed the particles. The scalebar indicates 50 µm. **(b**) The effect of 48 h 1, 10 or 100 µg/mL rutile treatment on the viability (MTT; blue) and cytotoxicity (LDH; orange) of microglial cultures. For the MTT measurement, relative viability, for the LDH measurement, cytotoxicity is given as a percentage of control, solvent-treated values. Data are from one experiment, with 2-6 parallel measurements in each concentration range. Error bars represent SEM. Kruskal-Wallis: p<0.001: ***

As these results suggest that microglial cells may play a primary role in the phagocytosis of nanoparticles that reach the brain, we also investigated the effects of anatase and rutile nanoparticles on this cell type. For this purpose, cortical glia cultures from new-born mice were prepared, which showed patches of microglial cells on their surface after 1-2 weeks in culture. Spherical microglia cells with fine processes were isolated from the surface of the flat astroglia cells by gentle washing with medium and subsequently replated on tissue culture plates. Since 48-hour treatment with rutile had shown the most pronounced effect in cortical astroglia cultures, we investigated the effects of 48-hour rutile exposure at concentrations of 1, 10, 50, and 100 µg/mL using both MTT and LDH assays. Rutile nanodispersion decreased cell viability (one-way ANOVA, p=0.010) and increased cytotoxicity (one-way ANOVA, p=0.015) in microglial cells after 48 hours of treatment (**Figure 6b**), with a statistically significant increase in cytotoxicity observed only at the highest concentration of 100 µg/mL.

## 4. Discussion

Due to its favorable properties, titanium dioxide is widely used in industry, mainly in the form of nanoparticles ranging in size from 1 to 100 nm. Because of its whitening characteristics, it is also known as “titanium-white” and is commonly used in the paint and cosmetics industries (Haider et al., 2019). TiO_2_ has also been utilized as a food additive, as an early safety assessment conducted by the Joint FAO/WHO Expert Committee on Food Additives (JECFA) in 1969 concluded that it was harmless: “Titanium dioxide is a very insoluble compound. The studies in several species, including man, show neither significant absorption nor tissue storage following ingestion of TiO_2_. Establishment of an acceptable daily intake for man is considered unnecessary” (JECFA, 1970). However, more recent studies have shown that TiO_2_ nanoparticles may pose health risks, particularly with prolonged exposure (Z. Chen et al., 2020). These findings led the European Union to ban the use of TiO₂ (E171) as a food additive starting from January 2022 (European Commission, 2022). Nevertheless, its use remains widespread in other sectors.

The neurotoxicity of nanoparticles has been demonstrated in numerous studies (Gong et al., 2022). However, the exact mechanisms involved, as well as the degree to which different cell types within the complex neural tissue are affected, remain incompletely understood (Ge et al., 2019). The application of an *in vitro* system may help address these gaps, as they are widely used in various fields of toxicological research (Madorran et al., 2020). In case of titanium dioxide, previous experiments were primarily conducted on human or rat cell lines, although primary cultures have also been used but to a lesser extent (Wang et al., 2022). In most cases, concentration- and exposure-dependent cytotoxic effects, namely a reduction in cell viability, have been reported (Hong et al., 2015; Papp et al., 2020). Mechanistically, these toxic effects are often associated with the generation of excessive reactive oxygen species (ROS), leading to oxidative stress, lipid peroxidation, and DNA damage (Ferraro et al., 2020; Huerta-García et al., 2014; Ze et al., 2013). Additionally, TiO₂ nanoparticles have been shown to impair mitochondrial function, disrupt calcium homeostasis, and activate pro-inflammatory signaling pathways such as NF-κB, resulting in the release of cytokines and chemokines (Han et al., 2013; Long et al., 2007; Sheng et al., 2015). These responses are not only cytotoxic but may also impair neuronal signaling and glial support functions, potentially contributing to broader neurodegenerative processes (Hsiao et al., 2016).

It is increasingly recognized that the various cell types constituting the central nervous system differ not only in function, but also in their susceptibility to external insults, including nanoparticle-induced toxicity (Chang et al., 2021; Feng et al., 2015; Limón-Pacheco et al., 2020). While most of previous toxicological investigations have focused on neurons and have identified direct neurotoxic mechanisms, such as oxidative stress, mitochondrial impairment, and excitotoxicity (Gerber et al., 2022; He et al., 2018; Hong et al., 2015; Mu et al., 2020; Sheng et al., 2015; Valentini et al., 2018; Wu et al., 2010; Wu & Xie, 2016), glial cells have received comparatively less attention (Coccini et al., 2015; De Simone et al., 2016; Papp et al., 2020; Pérez- Arizti et al., 2020; Wilson et al., 2015). Importantly, during the last decades it was proven that astrocytes are not merely passive supporting cells within the CNS but active participants in both physiological and pathological processes (Verkhratsky & Nedergaard, 2018). They fulfill diverse roles including regulation of the extracellular ionic and metabolic environment, synapse formation and maintenance, blood-brain barrier integrity, and modulation of neuro-immune responses. Their previously assumed homogeneity has been challenged by recent studies showing significant regional heterogeneity, with distinct gene expression profiles (Batiuk et al., 2020; Endo et al., 2022) and functional characteristics (Xu et al., 2001; Zhao & Flavin, 2000) that may influence their vulnerability to toxic insults such as oxidative stress. Similarly, microglia is also known to respond rapidly to oxidative and inflammatory stimuli (Ishihara & Itoh, 2023; Rojo et al., 2014), and may act as amplifiers of nanoparticle-induced damage through the release of pro-inflammatory mediators (Huang et al., 2022; Long et al., 2007; Xue et al., 2012). These considerations highlight the importance of evaluating neurotoxic effects not only at the neuronal level, but within the broader context of neuron–glia interactions, which may significantly influence the outcome of toxic exposure in complex neural systems.

In industrial applications, titanium dioxide nanoparticles are predominantly produced in two crystal forms: rutile, which is typically rod-shaped, and anatase, which is generally spherical (Allen et al., 2018). Furthermore, much of the research utilizes only one form, preferably anatase (Coccini et al., 2015; He et al., 2018; Hong et al., 2015; Mu et al., 2020; Sheng et al., 2015; Valentini et al., 2018) or a non-defined mixture of both crystal forms (Long et al., 2007; Valdiglesias et al., 2013; Wilson et al., 2015). To the best of our knowledge, only one study has compared the effects of anatase and rutile TiO₂ nanoparticles on neural cells (Wu et al., 2010), mostly using a cell line (PC12) that does not fully replicate the properties of native neural cells. Therefore, we systematically applied a comparative analysis on the effects of anatase and rutile nanoparticles on different neural cell types, both in pure and in mixed-culture models.

### 4.1. Effects of anatase on different cell types

In our experiments, the anatase form was applied at different concentrations to different types of primary cultures during 24 and 48 h treatments. Although some studies suggest that the anatase form is the most toxic (Baranowska-Wójcik et al., 2020), our results do not support this observation.

Anatase did not cause a significant reduction in cell viability in any of the culture types (hippocampal and cortical astroglia cultures, neuron and cortical glia containing mixed culture) after 24 h of treatment, neither in MTT nor in LDH measurements. An effect was observed only in cortical astroglia cells after 48 hours, and this was detectable solely by LDH measurements. These results are in contrast with studies reporting significant viability loss following anatase exposure, although such effects in astrocytes were typically observed only at very high (>600 µg/mL) concentrations (Papp et al., 2020). It is important to note that comparability between studies is greatly complicated by the use of different cell models. Most studies reporting toxicity thresholds in the range of 10–60 µg/mL utilized primary neurons derived from the hippocampus (He et al., 2018; Hong et al., 2015; Mu et al., 2020; Sheng et al., 2015) or used cell lines (Coccini et al., 2015; Hsiao et al., 2016; Long et al., 2007; Valdiglesias et al., 2013; Wu et al., 2010).

Another interesting observation was that anatase increased the relative cell viability in mixed cultures. One possible explanation for this can be that titanium dioxide causes oxidative stress in the cells. One possible mechanism for this is a “ROS-induced ROS release”, whereby reactive oxygen species (ROS) act on the inner membrane of mitochondria, modifying the function of anion channels, leading to further ROS release. The accumulated ROS, in turn, leads to mitochondrial and cellular damage (Huerta-García et al., 2014). It is possible that oxidative stress increases the reduction capacity of neurons which can be detected as an increased cell viability by the MTT method. However, the exact mechanism behind that can only be revealed by further studies, e.g. by examining changes in mitochondria. Since we observed a similar effect in case of the rutile form, the phenomenon seems to be independent of the type of crystal structure. It is also important to note that the increased viability detected at higher concentrations of NPs can be considered as an artifact due to the “nano snow” layer formed on the cells above 100 µg/mL of NPs. In these cases, the actual NP concentration reaching the cells was probably much lower than the nominal concentrations, due to aggregation.

The different types of cellular responses to anatase and rutile may be based on the different crystal structures of the nanoparticles. Jin et al. (Jin et al., 2011) found that the two forms have different effects on ROS generation. After dispersion, there will be more reactive hydroxyl groups on the surface of the NPs, which can lead to the formation of superoxide and hydroxyl radicals by electron capture. The study also showed that the anatase form is more effective to form radicals due to its larger surface area than the rutile form. Both forms were detectable in the cytoplasm of the HaCaT cells used in that study, but the anatase form was also located in the nucleus and thereby triggered genotoxicity. The anatase form was also found in the mitochondria, where it could influence the function of the electron transport chain, resulting in overproduction of reactive oxygen species. Unlike anatase, rutile was not identified in the nucleus or the mitochondria. Nevertheless, the authors did not exclude the possibility that rutile may also be able to enter into these structures and damage them (Jin et al., 2011). All of this suggests that the different origins, purity and condition (e.g. crystal structure, crystal size) of the nanoparticles may be responsible for the different results.

It may be questioned to what extent the observed effect depends on the presence of nanoparticles and to what extent on the specific effect of chemical components (e.g. ions). Titanium dioxide is practically insoluble in water, so the ionic effect of titanium can be excluded in our studies.

### 4.2. TiO_2_ with rutile crystal structure acts differently on different cell types

In the literature, data on the effects of rutile on cells are mostly limited to studies in which it is used in combination with anatase (Long et al., 2007; Valdiglesias et al., 2013; Wilson et al., 2015). To our knowledge, no results have been published on the specific effects of rutile on neurons or astrocytes, despite rutile being the predominant form produced in industrial applications.

In our study, rutile was applied to various cell cultures at different concentrations and treatment durations (24 h or 48 h). Similarly to anatase, an increase in cell viability was observed in mixed cultures containing both neurons and glial cells. However, when rutile was applied to cortical astroglia cultures, a significant reduction in relative cell viability (MTT) and an increase in cytotoxicity (LDH) were observed, in a time-dependent manner. Effects in hippocampal glia cultures were only detected at the highest concentration tested (1000 µg/mL). These findings suggest that glial cells, unlike neurons, are more sensitive to rutile exposure.

One possible explanation for the absence of detectable toxicity in mixed cultures is the predominance of neurons over glial cells in this culture type. If neurons are not sensitive to titanium dioxide exposure, a reduction in glial cell viability might still occur but be masked by the stronger viability signal from the neurons.

Our bioinformatic analysis revealed that, beyond the previously identified general transcriptomic profile (Endo et al., 2022), hippocampal and cortical astrocytes also differ in the expression of oxidative stress-related genes. These differences may influence the cells’ susceptibility and responsiveness to oxidative stress. By applying curated gene sets (Table A1), associated with oxidative stress, glutathione metabolism, superoxide metabolism, and NAD(H) homeostasis, we identified six genes that displayed differential expressions between cortical and hippocampal astrocytes. Specifically, *C4b* (Complement component 4B involved in complement pathway activity), *Cyba* (Cytochrome b-245 alpha chain, a part of NADPH-oxidase complex), *Agt* (Angiotensinogen, involved in neuroinflammatoric responses), and *Gstm6* were upregulated, while *Gpd1* and *Ipcef1* were downregulated in hippocampal astroglia. The increased expression of *C4b*, *Cyba*, and *Agt* suggests a pro-inflammatory and oxidative environment, whereas the elevated levels of *Gstm6* (Glutathione S-transferase Mu 6) may contribute to a more robust homeostatic and counterbalancing antioxidant defense. In contrast, the reduced expression of *Gpd1* (Glycerol-3-phosphate dehydrogenase 1, involved in redox metabolism) and *Ipcef1* (Interaction protein for cytohesin exchange factors 1, involved in cytoskeletal and migratory dynamics) may underlie a shift toward a hyporeactive astroglial phenotype, potentially diminishing responsiveness to neuroinflammatory cues. Although we did not directly assess the transcriptomic landscape of astroglial cells under *in vitro* conditions, astrocytes can maintain certain region-specific molecular signatures even outside the *in vivo* context (Cragnolini et al., 2018; Ernsberger et al., 1990; Kipp et al., 2008).

Since the cortical astroglia cultures showed reduced viability in response to rutile, we also treated pure microglial cultures with rutile for 48 hours at concentrations of 1, 10, 50, and 100 µg/mL. In microglial cells, a tendency toward reduced viability was observed using the MTT assay, while the LDH assay revealed a significant increase in cytotoxicity only at the highest concentration (100 µg/mL). These findings - especially in the context of rutile - are intriguing and novel observations. Former studies have investigated the effects of TiO₂ nanoparticles using the BV2 cell line, but typically with either anatase or a mixture of crystal forms. BV2 cells have also been shown to be sensitive to TiO₂ exposure, with toxicity thresholds as low as 5–10 µg/mL. In agreement to our findings, internalization (phagocytosis) of nanoparticles by BV2 cells has been also reported (Hsiao et al., 2016; Long et al., 2007; Valdiglesias et al., 2013).

This study highlights the glial cell specific effects of anatase and rutile crystal forms TiO₂, nanoparticels on various neural cultures. Further investigations, particularly focusing on mitochondrial function and ROS generation, are needed to better understand the underlying pathways of TiO₂ induced cellular responses.

## 5. Conclusions

Taken together, substances containing nano-sized titanium particles in the rutile form may pose a specific risk to key cell types in the central nervous system, particularly to astroglial cells. Impairment of astrocyte function can result in widespread neurophysiological disturbances and may indirectly compromise neuronal viability and synaptic integrity. Although the rutile form of TiO₂ is generally considered less reactive than the anatase form, emerging evidence suggests it can still elicit adverse effects under certain conditions. These observations underscore the importance of conducting more cell-type-specific investigations to better understand astrocyte vulnerability and their potential role in nanoparticle-induced neurotoxicity.

## Abbreviations

TiO_2_: titanium dioxide
MTT: 3-(4,5-dimethylthiazol-2-yl)-2,5-diphenyl tetrazolium bromide
LDH: lactate dehydrogenase

## Supplementary Table S1

The following oxidative stress-related gene sets from GSEA Molecular Signature Database (https://www.gsea-msigdb.org) are used to filter the differentially expressed genes (false discovery rate (FDR) < 0.05 and absolute log2 fold change (|log2FC|) > 1) between hippocampal and cortical astroglia data set (GSE198024).

**Table S1.** Oxidative stress-related Molecular Signature gene sets (from GSEA) used for the analyses.

| Name of Gene Set | Version | ID |
| --- | --- | --- |
| Hallmark Reactive Oxygen Species Pathway | v2025.1.Mm | MM3895 |
| WP Oxidative stress and redox pathway | v2025.1.Mm | MM15823 |
| WP Oxidative stress response | v2025.1.Mm | MM15941 |
| WP Oxidative damage response | v2025.1.Mm | MM15945 |
| GOCC NADPH Oxidase Complex | v2025.1.Mm | MM12311 |
| GOBP Superoxide Metabolic Process | v2025.1.Mm | MM4854 |
| GOBP Regulation of Superoxide Metabolic Process | v2025.1.Mm | MM10142 |
| GOBP NADH Metabolic Process | v2025.1.Mm | MM4836 |

**Figure S1.**
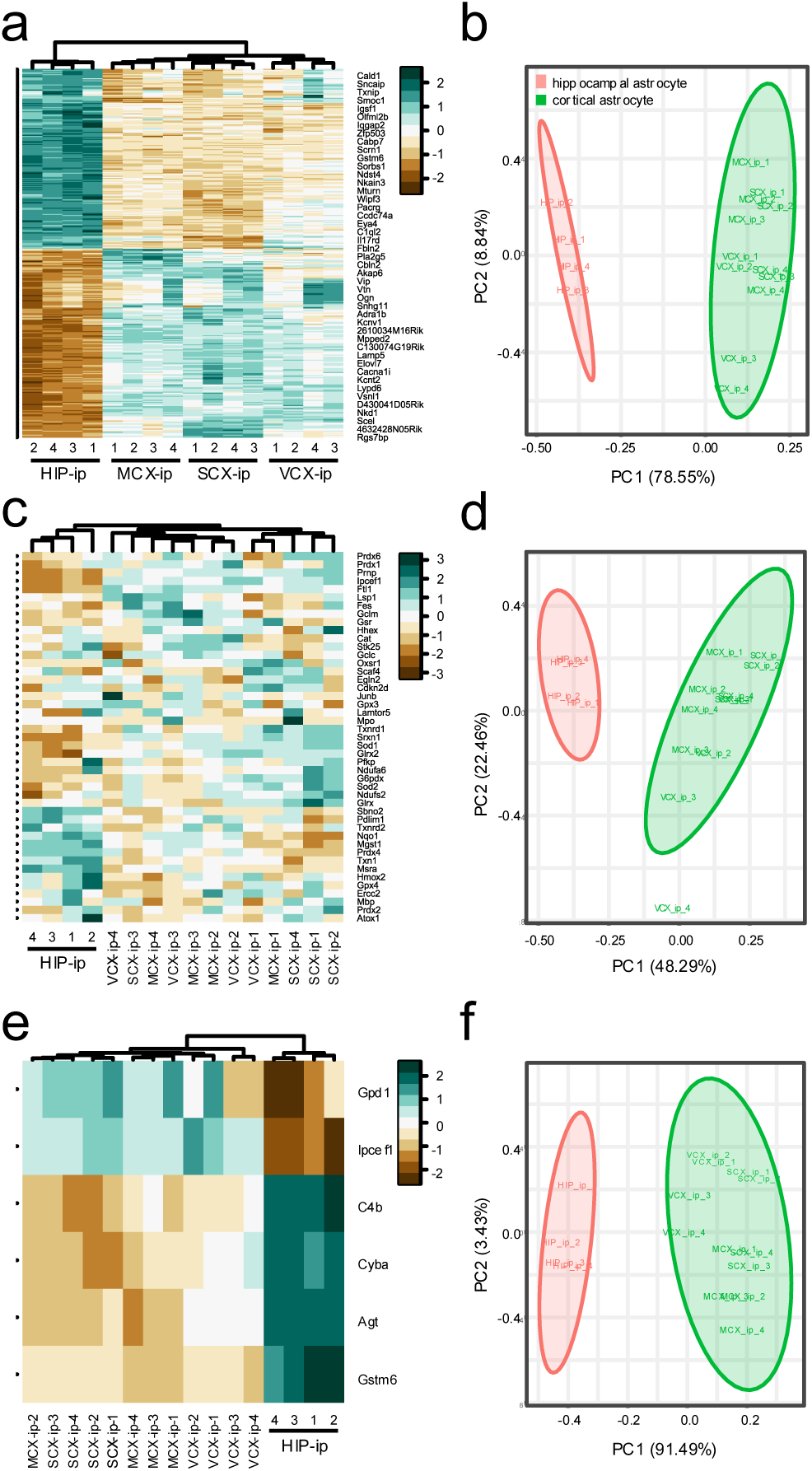
Hippocampal and cortical astrocytes differ in their general and oxidative stress transcriptomic profile. (**a**) Heat map showing the log_2_ fold change (FC) values of 359 differentially expressed genes (DEGs) between hippocampal (HIP) and cortical (from motor, sensory, and visual cortex - MCX, SCX and VCX, respectively) origins, with a threshold of log2 FC > 1, FDR < 0.05. “ip- #” represent the biological replicates in the original GSE198024 dataset (Endo et al., 2022). According to the dendrogram, astrocytes can be classified into 2 broad groups which is confirmed by the (**b**) PCA analysis. (**c**) Heat map showing the log_2_ expression values of 45 expressed genes from the Hallmark Reactive Oxygen Species Pathway GSEA gene set (MM3895, GSEA) in the same hippocampal (HIP) and cortical (MCX, SCX and VCX) samples. The dendrogram again shows that astrocytes cluster into two broad groups, which is confirmed by the (**d**) PCA analysis. (**e**) Heat map showing the log_2_ fold change (FC) values of six differentially expressed genes (DEGs) between hippocampal (HIP) and cortical (MCX, SCX and VCX) samples filtered using the DEGS with Oxidative Stress and redox pathway-related gene sets (see Table S1) with a threshold of log2 FC > 1, FDR < 0.05. The dendrogram shows the classification of astrocytes could be classified into 2 broad groups, confirmed by the (**f**) PCA analysis.

